## Supplementary figures and images for "Post-Insemination Selection Dominates Pre-Insemination Selection in Driving Rapid Evolution of Male Competitive Ability"

### Supplemental Figures

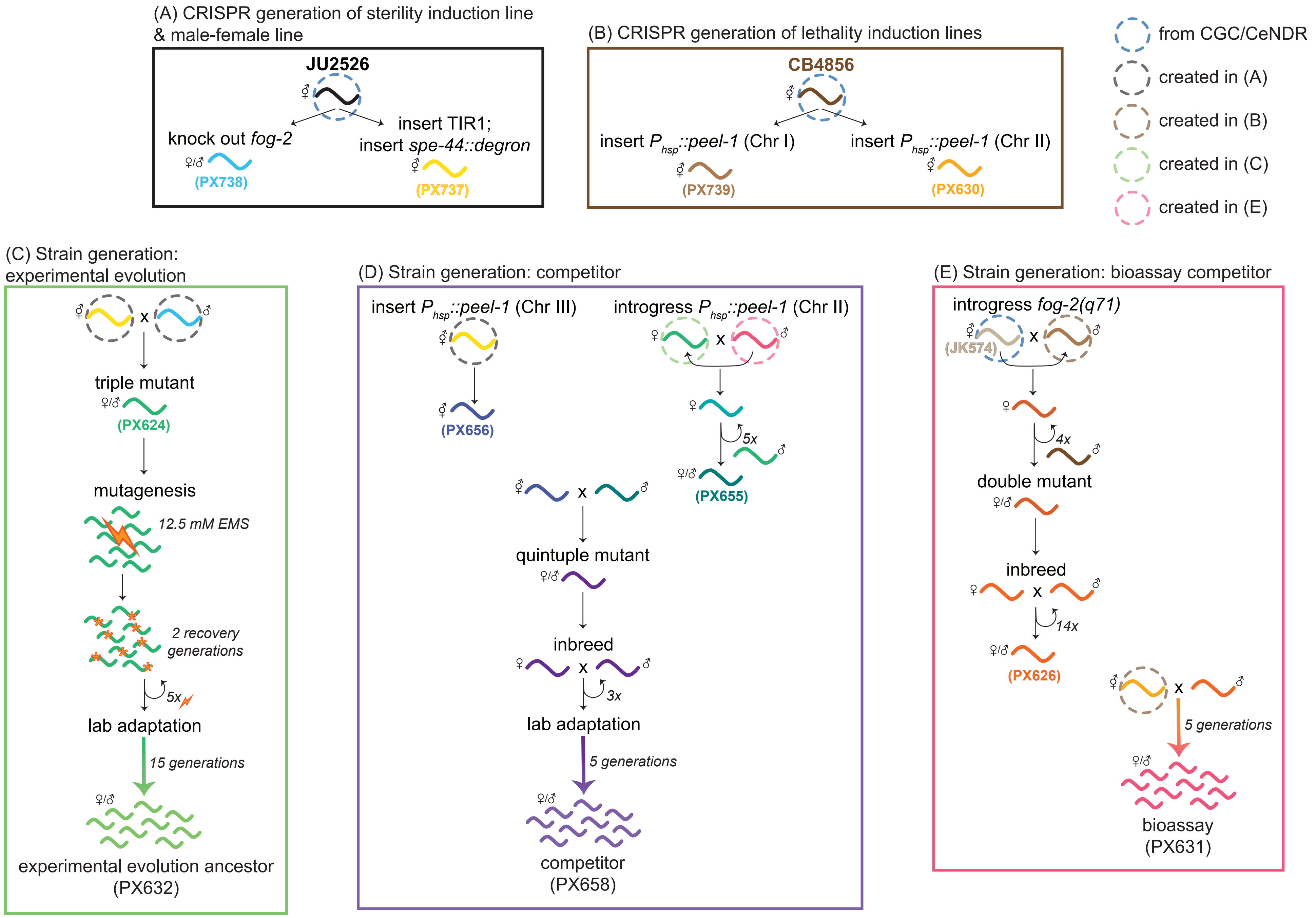

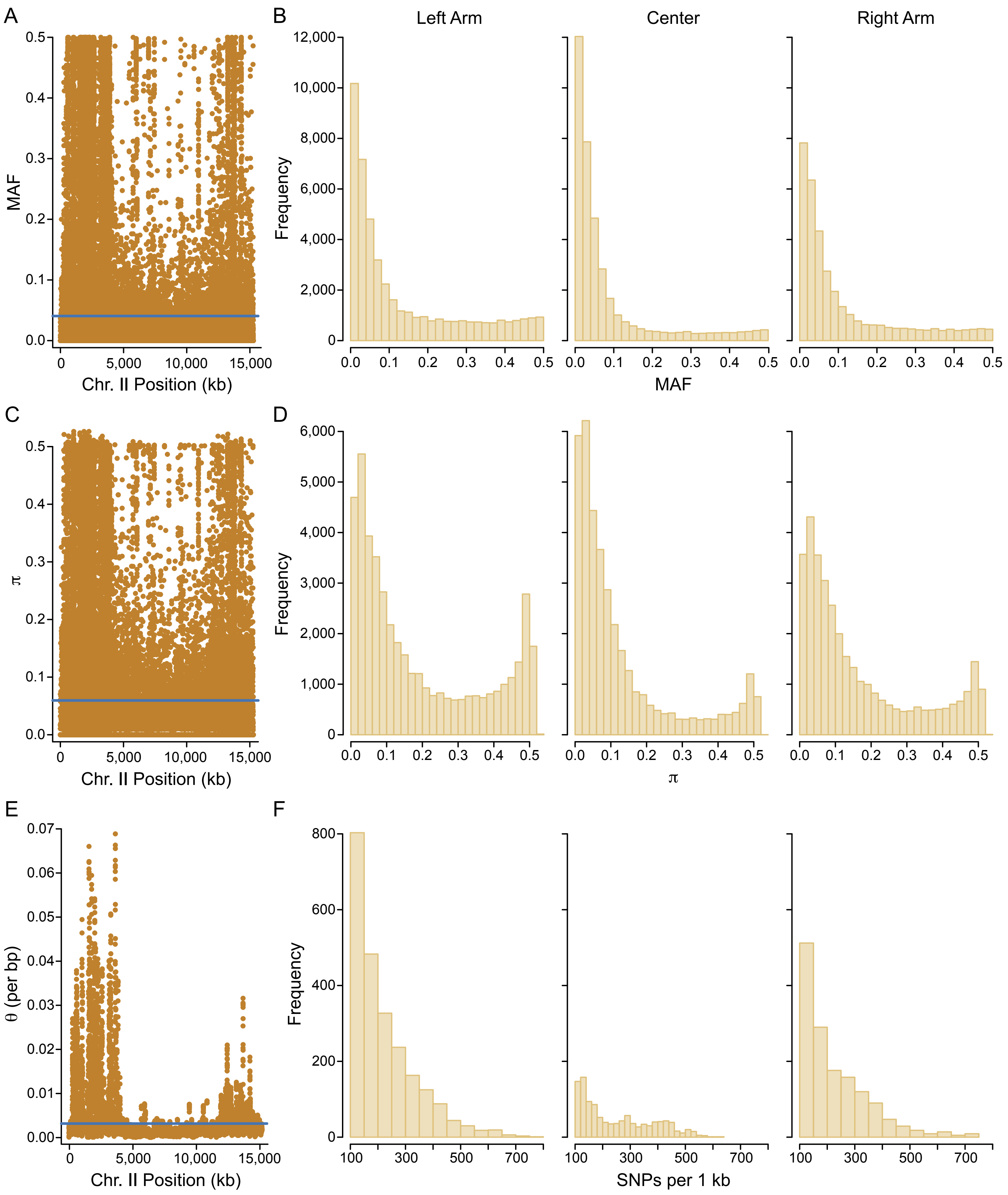

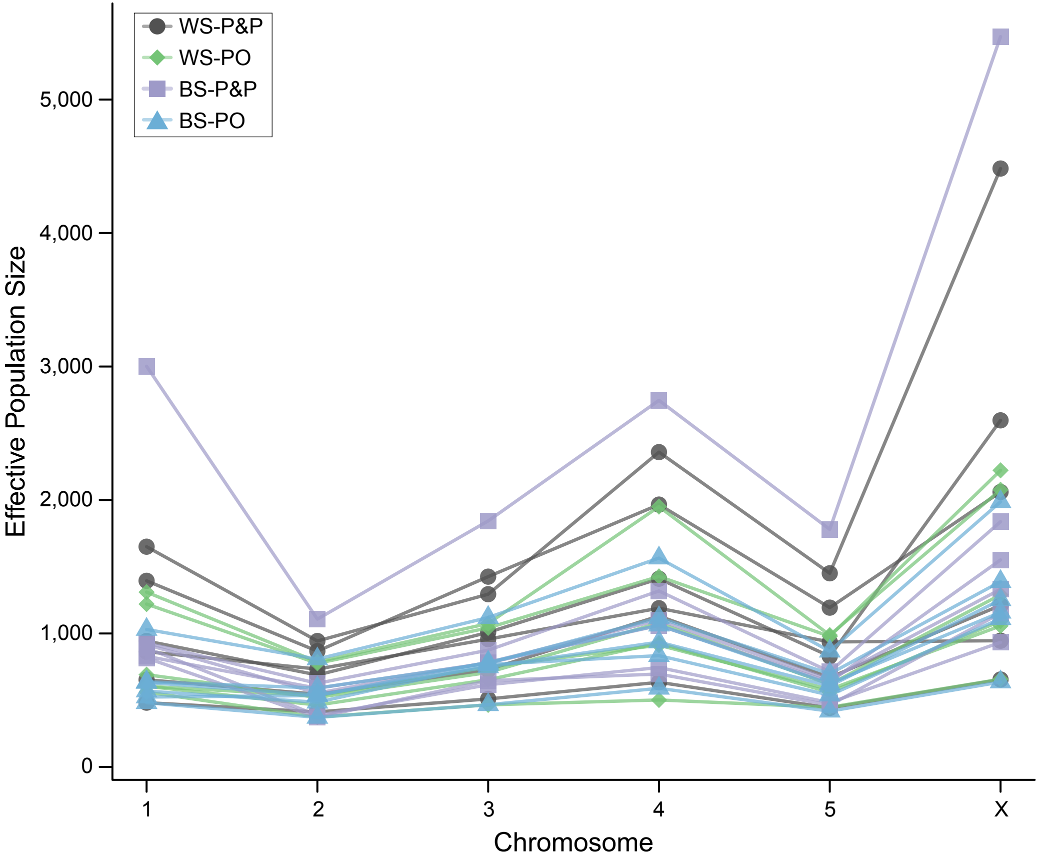

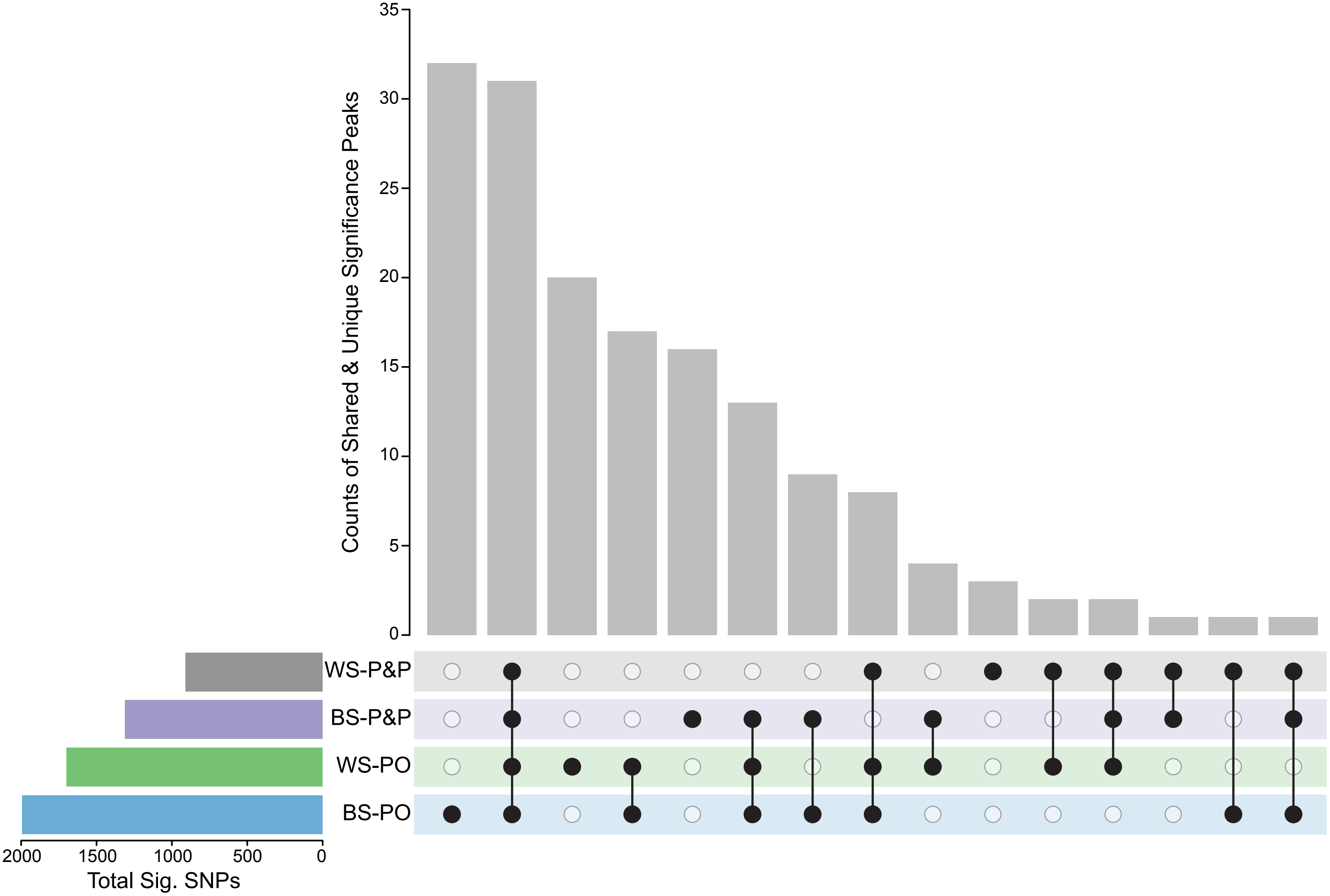

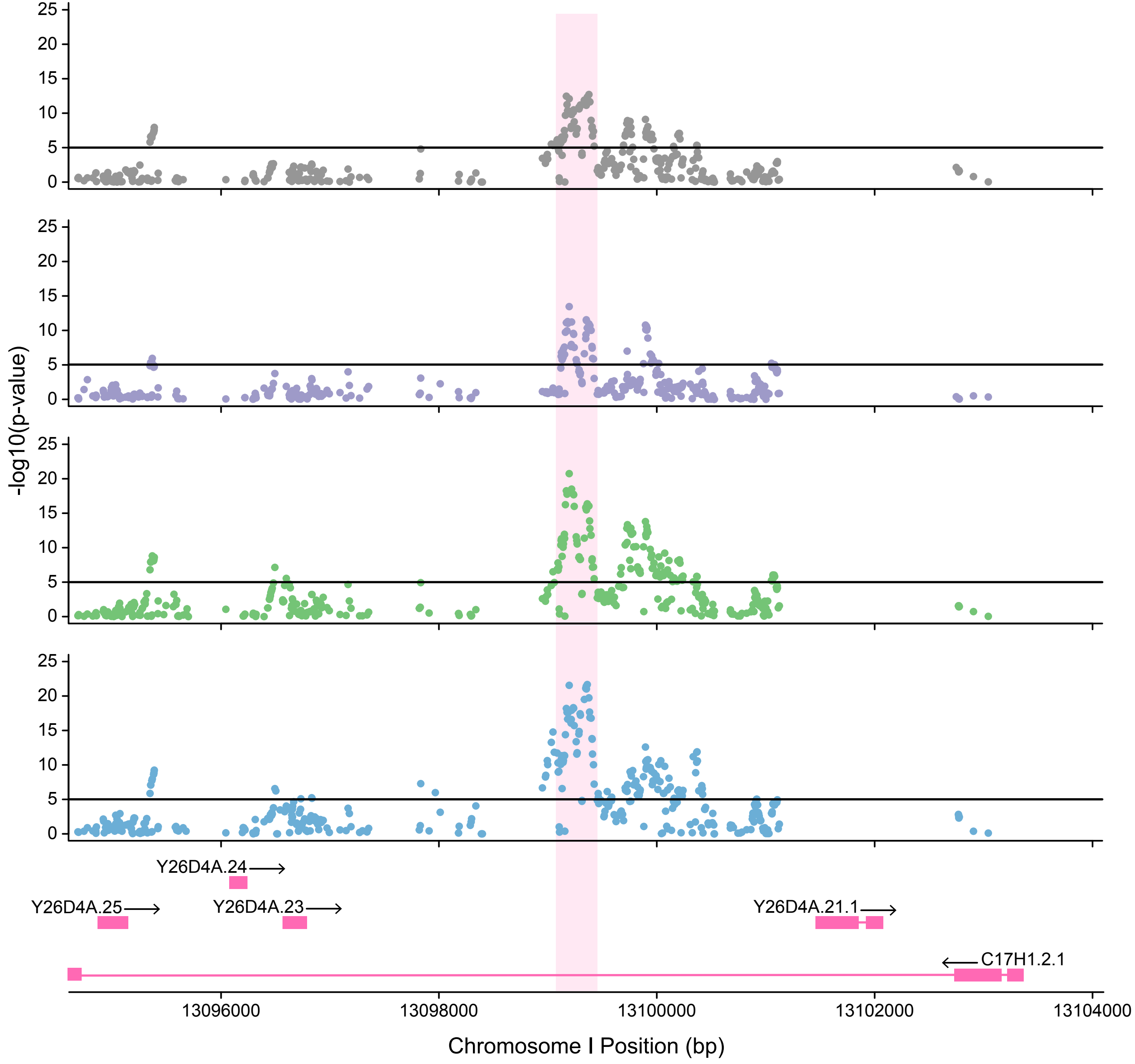

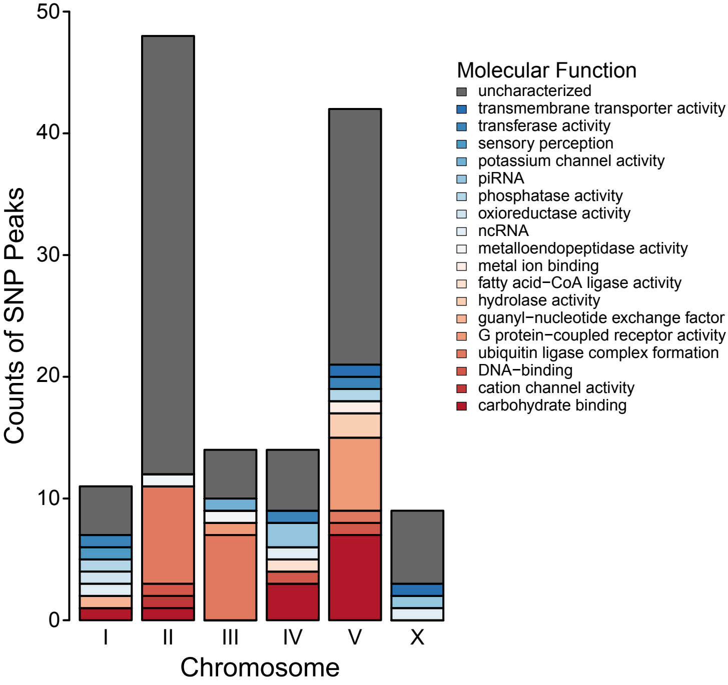
